# Exploring Dance Movement Therapy as a Novel Approach to Improving Heart Rate Variability in Healthy Older Adults

**DOI:** 10.64898/2025.12.14.694193

**Authors:** Laura Sebastiani, Said Daoudagh, Giacomo Ignesti, Marina Raglianti, Paolo Paradisi

## Abstract

A decline in cardiac autonomic control, as assessed by heart rate variability (HRV), is considered a natural component of ageing. However, physical exercise and dancing can be effective in preventing/reversing this decline.

Aim of this study was to verify whether Dance Movement Therapy (DMT), which uses dance, movement, body awareness and embodied interpersonal communication to promote well-being, improves vagally-mediated HRV, which has been linked to physical and mental health, in a sample of people aged over 65. To this purpose, HRV-derived vagal and sympathetic indices, obtained by electrocardiogram signals, as well as tonic skin conductance level, which is a marker of tonic sympathetic activity, recorded during two rest sessions that preceded and followed a 8-week DMT activity, were compared through repeated measures analysis of variance.

Results showed post-DMT changes in HRV-derived vagal parameters in the time, frequency and nonlinear domains. More specifically, the root mean square of the successive differences between RR intervals (RMSSD), the high frequency power (HF) and the standard deviation of the beat-to-beat changes in RR intervals (short-term HRV) (SD1) were higher in the post-than pre-DMT session. Conversely, no changes were found in HRV sympathetic indices nor in skin conductance level.

The post-DMT improvement in vagally-mediated HRV in healthy old people is a new finding and we argue that this beneficial effect may be due to DMT’s ability to enhance social interaction and promote well-being.

## 1. Introduction

Heart rate variability (HRV), which is defined as the variation in the time intervals between successive heartbeats (RR), is widely used by the scientific community to indirectly investigate the dynamics of the autonomic system based on electrocardiogram measurements [1, 2].

The study of HRV employs a variety of non-invasive metrics, including statistical indices that span from time averages and frequency-based analyses to nonlinear, or complex, indices, which reflect the continuous and dynamic interplay between two branches of the autonomic nervous system: the sympathetic nervous system (SNS), which prepares the body for action, and the parasympathetic nervous system (PSNS), which promotes rest and recovery [2]. A high HRV is associated with particularly strong PSNS activity, which is mediated by the vagus nerve, and indicates good recovery, general physical fitness and well-balanced autonomic control. Conversely, low HRV reflects increased SNS activity and/or vagal withdrawal, and is indicative of the body’s response to stress, illness or overtraining [1].

HRV has also proved useful for investigating mental health. Actually, enhanced cognitive functions have been linked to high, vagally mediated, HRV at rest (e.g. mental flexibility, working memory, attention regulation, motor imagery) [3, 4, 5]. In contrast, low resting state HRV values have been linked to cardiovascular disturbances, as well as to impaired adaptability to both internal and external stimuli [6].

Within the context of the ageing process, a reduction in HRV indices (i.e. reduction in RMSSD, High frequency (HF) and Low Frequency (LF) power) is widely accepted as a natural component of the ageing human condition [7], and lower cardiac vagal control in older people has been found to be associated with cognitive impairment and functional decline [8].

However, many studies that described the age-related changes in HRV have focused on older people with medical problems. For instance, older adults with clinical depressions showed low LF-HRV that is indicative of reduced baroreflex function [9]. Also, a longitudinal analysis involving older women with late-onset depressive symptoms showed a significant enhancement of LF/HF balance with time, indicative of a progressive long-term increase of sympathetic tone. [10].

Interestingly, a study by Tan and colleagues (2019) [11], found no age-related decline in HRV among older participants who had very few physical or psychological symptoms. Similarly, a study assessing the influence of age, gender, body mass index, and functional capacity on HRV in a cohort of healthy persons [12], showed that parasympathetic HRV indexes (i.e. RMSSD, HF) exhibited a rapid decrease until the fourth decade of life, and subsequently remained relatively stable in older age groups.

Therefore, when assessing alterations of cardiac autonomic control in older individuals, other factors besides old age must be taken into consideration. For instance, evidence has been presented which demonstrate that lifestyle factors, such as physical activity levels, can influence the body’s normal functioning and modulate HRV [13]. Indeed, the tendency towards sedentary behaviour that is frequently exhibited by the older population has been linked to an elevated prevalence of cardiovascular diseases [14]. Conversely, the efficacy of aerobic exercise in enhancing metabolic and cardiovascular health in older adults has been demonstrated [15], thereby indicating that effective strategies can be employed to prevent or reverse HRV deterioration in this demographic. In particular, dancing represents a form of aerobic exercise with which older adults can easily engage and research has demonstrated that dancing can yield a number of benefits for this demographic [16].

More specifically, the physical advantages obtained were comparable to those achieved through physical exercise training, including improvements in flexibility and postural stability [17]. Furthermore, a positive impact has been observed in the domain of psychosocial well-being and in sleep quality. [18, 19]. Recently, it has been demonstrated that an exergaming-based dance training protocol can improve heart rate variability (HRV) in healthy older adults [20]. Specifically, after a six week training, participants showed an improvement in HF power, RMSSD and pNN50 compared to the control group.

In this regard, Dance Movement Therapy (DMT) is a particularly noteworthy approach as it emphasises the integration of emotional, physical and social dimensions. In fact, DMT^1^ is a complementary therapy that promotes emotional, cognitive, physical and social integration through interactions between individuals mediated by body movements [24, 25]. DMT emphasises the expressive and qualitative dimensions of movement as vehicles for improving psychological and emotional well-being [26] and is based on the assumption of a dynamic relationship between mind, body and social interactions, which includes both affective and cognitive processes. It is rooted in the theoretical framework of *embodied cognition*, which posits that cognitive processes are intrinsically shaped and modulated by our body’s engagement with the surrounding environment, including the social environment [27, 26, 28].

DMT activity has demonstrated efficacy in enhancing resilience, as well as mood and relaxation, in chronic pain patients [29] and in improving distress and anxiety in cancer patients [30]. Incorporating DMT elements into physical exercises has also been effective in enhancing the fitness and overall functioning of older nursing home patients in wheelchairs [31] and improving mood of older people in acute hospital settings [32]. Furthermore, it has been effective in enhancing general well-being and perceived quality of life of healthy population including the olders [33].

The objective of the present study was to verify the potential effectiveness of Dance Movement Therapy (DMT) in improving HRV in a sample of Italian individuals over the age of 65. To this end, we implemented a one-group pretest–posttest design, which is a quasi-experimental research method often used by behavioural researchers and social scientists to evaluate the potentialities of an activity on a specific sample [34]. Our working hypothesis is grounded, by one hand, on preceding findings demonstrating a correlation between elevated HRV and enhanced physical and mental health and, on the other hand, on evidences about effectiveness of DMT in promoting well-being among older people. We then hypothesised that participation in the DMT activity may yield favourable outcomes in terms of enhancing vagal-mediated HRV within the present sample of over-65 individuals. A possible DMT-induced reduction in skin conductance tonic level (SCL) was also expected, since this is considered a marker of tonic sympathetic activity and a useful indicator of stress levels, with higher SCL values generally correlating with higher levels of arousal and stress. Furthermore, since poor sleep quality and sleep deprivation have been found to be associated with reduced HRV [35] and changes in sleep naturally occur with age (e.g. decreased sleep efficiency and more frequent nocturnal awakenings), a secondary objective of the present study was to verify whether sleep quality could influence the HRV of our sample of older people and whether the DMT activity could modulate not only sleep quality *per se* but also the possible association between HRV and sleep quality.

Finally, since older people are generally at greater risk of depression than younger people [36], often because of their living conditions, and depression severity is negatively associated with HRV, a further aim was to investigate whether the presence of depressive symptoms in our sample of older people could influence HRV and whether the participation in DMT activity could reduce depression symptoms and modulate the possible HRV-depression association.

In the next Section 2 we explain in detail the tools that we applied for the statistical analyses. In Section 3 we report the results of the statistical analyses, while in the Section 4 we discuss our results, giving some interpretation also in the view of previous literature. For reader’s convenience, in Appendix A we also report a list of the acronyms used in the text.

## 2. Methods

### 2.1. Experimental design and participants

Recruitment advertisements were prepared and disseminated via flyers that were distributed at territorial associations and gyms, and also through local newspaper channels. People who answered the advert were interviewed by PP or SD to collect some general personal information (like age, sex^2^, previous job, general health, and how much exercise they did before and now). This information was put into a database to help choose the right people to partake in the study. Participants of both sex were included in the study if they met the following criteria: they were over 65 of age; they were not affected by neurological (e.g., Parkinson’s disease), psychiatric (e.g. social phobia), musculoskeletal (e.g. arthritis), cardiovascular (e.g. hyperthension) disorders; they were able to carry out their daily activities independently; they reported participating in leisure motor activities to a moderate extent, such as walking at a brisk pace, cycling on flat terrain, gym activities (e.g., pilates), ballroom dance, for approximately one hour, two or three times a week; they have a body mass index (BMI) in the range of 23 − 29 kg/m², which is recommended for people over the age of 65; they do not drink coffee regularly or make moderate use of it (no more than two coffees a day).

A total of 27 persons (females, *N* = 17) were found to be eligible and enrolled in the study. Participants took part in a eight-week DMT cycle on a once-weekly basis. All the DMT meetings of the cycle were scheduled at the same time of the day, namely from 9:30 to 10:30 a.m.. The DMT cycle was preceded (pre-DMT) and followed (post-DMT) by a one-hour session where participants answered the self-report questionnaires Geriatric Depression Scale (GDS) and Pittsburg Quality of Sleep Index (PSQI), and resting state physiological parameters (ECG and GSR) were collected. The pre-DMT and post-DMT sessions were done within one week before and after DMT activity, respectively.

Two DMT cycles were scheduled over the course of two successive periods of the year: Cycle 1 from January to March and Cycle 2 from March to May. Participants were assigned to one cycle or the other in order to obtain two groups (1st and 2nd cycle) of similar size (1st cycle, n=13, and 2nd cycle, n=14). and with a similar female-to-male ratio (1st cycle, 9 F and 4 M; 2nd cycle, 9 F and 5 M). The decision to divide the participants into two groups was taken in accordance with the recommendation of the DMT professional (MR) who conducted the DMT sessions. This professional indicated that a sample size of 14 would be optimal for facilitating the development of group dynamics. The DMT professional registered participants’ attendance at the DMT cycle at each of the eight DMT meetings. Only those who attended at least six out of eight meetings and did not report any motor disturbances throughout the cycle were included in the subsequent analysis. However, one male participant from the 1st cycle attended only 3 of the 8 DMT sessions and was therefore not included in the subsequent analysis. One male participant from the second cycle was excluded from the analysis due to a minor bicycle accident that impaired his movement during subsequent DMT sessions. None of the other participants reported relevant events outside their daily routine. It is noteworthy that the remaining participants attended a minimum of 85% of the DMT sessions, with no reported instances of motor disturbances throughout the duration of the DMT cycle. Thus, the analysed sample comprised two groups: 1st cycle, n=12, and 2nd cycle, n=13. Demographic data are presented in Table 1.

**Table 1:** Demographic characteristics of participants.

| **Cycle** | 1 | 2 |
| --- | --- | --- |
| **Sample size** | N=12 | N=13 |
| **Age (mean±SD)** | 68.25± 3.41 | 72.18±6.26 |
| **Sex** | % | % |
| *female* | 66.7 | 72.73 |
| *male* | 33.3 | 27.27 |
| **General Health (subjective evaluation)** | N | N |
| *fairly good* | 1 | 4 |
| *good* | 7 | 5 |
| *very good* | 3 | 4 |
| *excellent* | 1 | 0 |
| **Ongoing leisure motor activities** | N | N |
| *walks* | 10 | 11 |
| *cycling* | 6 | 7 |
| *dance* | 1 | 1 |
| *gym* | 1 | 2 |
| *swimming* | 0 | 1 |
| **Previous motor activities** | N | N |
| *yes* | 11 | 13 |
| *no* | 1 | 0 |

This study was performed in accordance with the Declaration of Helsinki ethical standards and approved by the Committee on Bioethics of the University of Pisa (resolution no. 29/2024 of 26/07/2024). All participants read and signed the informed consent to participate in the study.

### 2.2. Research Instruments and procedure

The pre-DMT and post-DMT experimental sessions were scheduled between 9:30 a.m and 3:00 p.m. In accordance with the instructions provided during the recruitment stage, participants were required to refrain from consuming coffee for a period of at least two hours prior to attending the experimental session. Moreover, it was recommended that participants scheduled for a session in the early afternoon should consume a light meal at lunch. Upon arrival, participants were informed of the experimental procedures and provided written informed consent. They were then invited to sit in a comfortable armchair and equipped with the g-Nautilus 32 system (gTech, Schiedlberg, Austria) [37] to record synchronized electroencephalogram (EEG) and ECG signals and the wireless wearable sensor Shimmer3 GSR+ [38] for galvanic skin response (GSR) recording. The recording session lasted about 13 minutes. For ECG recording, 2 Ag/AgCl disposable electrodes were placed on the chest, according to ECG lead I, and connected to the EEG headset. Signals were acquired by means of the g.Recorder software (sampling frequency 500 Hz, band pass filtering 0.1 − 100 Hz) and ECG was analysed by means of Kubios HRV Premium software [39, 40] in order to obtain the series of consecutive distances between successive R peaks of ECG (RR series) and Heart Rate variability (HRV) measures. The following HRV components in the time and frequency domains were analyzed: the RR intervals (to assess heart rate); the root mean square of the successive differences between RR intervals (RMSSD), primarily reflecting parasympathetic (vagal) nervous system activity; the Stress Index (SI), which is the square root of Baevsky’s stress index [41] and serves as an indicator of sympathetic activity; the high frequency (HF, 0.15 − 0.4 Hz) components of the power spectrum density of the RR interval series that primarily reflect vagal activity related to the respiratory sinus arrhytmia. We also analysed a non linear metric derived from Poincaré plots that is SD1, that represents short-term HRV and is related to parasympathetic activity [40]. The electrodermal activity was monitored using the wireless wearable sensor Shimmer3 GSR+ [38] (sampling rate 128 hz), which allows real-time monitoring of the galvanic skin response (GSR). GSR is measured by passing a small current through a pair of electrodes placed on the palmar surface of two fingers of the non dominant hand. The skin conductance level (SCL, microSiemens), which consists of the slowly varying base signal of the GSR (i.e. tonic GSR), was measured as the average of data collected in the same temporal window as the ECG. The ECG-derived and SCL data were subsequently averaged over the 13-minute recording interval. Thereafter, analysis was conducted on the mean values obtained.

#### 2.2.1. Questionnaires

After each the recording session participants were invited to compile the italian version of the Geriatric Depression scale (GDS) [42] and of the Pittsburgh Sleep Quality questionnaire (PSQI) [43, 44].

The GDS is a 30-item instrument designed to evaluate the emerging symptoms of depression in elderly individuals. The scale comprises a series of questions and statements that are devised to evaluate the mood, feelings and behaviour of older people. The assessment encompasses the patient’s perception of their daily activities, their general mood, their sense of worthlessness, and any symptoms related to memory and cognitive function. Responses are collected in a binary format, with users providing a “Yes” or “No” response to each question. The final score can range from 0 (no symptoms) to 30 (severe depression) and can be interpreted as follows: 0–9, no depression; 10–19, mild depression; 20–30, severe depression.

The PSQI (Pittsburgh Sleep Questionnaire) is a self-report questionnaire frequently implemented in the evaluation of sleep quality. The tool is regarded as valuable due to its multifaceted approach, incorporating both subjective experiences and objective parameters. The global PSQI score (range 0-21) is obtained by summing seven component scores (range 0-3). 3 indicates the greatest degree of dysfunction. Higher scores indicate poorer sleep quality, with scores over 5 suggesting substantial sleep difficulties.

### 2.3. DMT protocol

The DMT protocol consisted of eight weekly DMT meetings, each lasting about one hour, which took place in a standard gym facility with sufficient space for unrestrained movement. THE DMT professional (MR), associated with APID [23], planned the settings and conducted the DMT meetings. In the second cycle of DMT, the same settings and order as in the first cycle were applied. Each DMT meeting was centred around a specific thematic element (e.g., exploration of the space around the body; finding unusual ways to move). The theme was developed within various phases that typically involved individual, couples, and group motor dialogues, with or without the use of accessories (e.g., elastic ribbons, masks, veils). With very few exceptions, almost all phases were accompanied by appropriate music. A more detailed description of the DMT protocol is enclosed in the Supplementary Material (Supplementary-Material-DMT-Protocol).

### 2.4. Statistical analysis

#### 2.4.1. Preliminary Analysis

The normality of data distribution was evaluated for all HRV indices, skin conductance and questionnaires using the Kolmogorov-Smirnov test. As the data were found to be normally distributed, parametric statistical analyses were employed.

The ECG recording of two participant, one of Cycle 1 and one of Cycle 2, were excluded from analysis because of some interferences that affected the recording during the Pre-DMT session. Two other subjects in Cycle 2 were excluded because their heart rate was below 50 beats per minute (bradycardia). Thus, HRV analysis was conducted on a total of 21 participants (Cycle 1, n=11; Cycle 2, n=10). Similarly, due to technical problems, the galvanic response of 2 participants of Cycle 2 could not be adequately recorded. Therefore, SCL analysis was conducted on a total of 23 participants (Cycle 1, n=12; Cycle 2, n=11).

RR and SCL data of the participants of the two cycles were preliminarly compared by means of separate repeated measures ANOVA with DMT (pre-DMT, post-DMT) as within-subject factor and Cycle as between-subject factor (Cycle 1, Cycle 2). Results showed that RR scores of the Cycle 2 (mean ± SD, 948.47 ± 115.34) were significantly (F(1,19)=7.15, p=0.015, *η*2 = 0.273; power= 0.72) higher than the RR scores of Cycle 1 (mean ± SD, 813.74 ± 115.34). Nevertheless, a lack of significant DMT × cycle interaction was found (F(1,19)=0.154, p=0.699), thereby suggesting that the DMT activity did not exert differential influence upon the cardiac activity of participants of the two cycles.

Results showed that SCL scores from Cycle 2 (0.486 ± 0.285) were not different from those in Cycle 1 (0.513 ± 0.284). In fact, no significant Cycle effect (F(1,20)=0.051, p=0.823) nor DMT x Cycle interaction (F(1,21)=3.932, p=0.061) were found. Consequently, for the purpose of subsequent analysis, data of participants from the two cycles were aggregated.

#### 2.4.2. Data analysis

The mean scores of the following metrics, GDS, PSQI, HRV parasympathetic indices (RMSSD,HF, SD1), Stress Index (SI), and SCL, measured in the two sessions, were analysed by means of separated repeated measures analysis of variance (rmANOVA) with DMT (Pre-DMT and Post-DMT) as within-subjects factor, and Age as covariate.

The possible association between Age and HRV indices and SCL was studied by means of Pearson correlation analysis. A linear regression model was used to analyse whether GDS and PSQI were predictive of HRV indices.

For all statistical test significance was set at *p <* 0.05. Bonferroni correction for multiple comparisons was applied when necessary. The SPSS.15 [45] statistical package was used for all analyses.

## 3. Results

### 3.1. HRV and SCL results

The mean values (± standard deviation) of RR and HRV indices relative to the pre- and post-DMT sessions are reported in Table 2. Repeated measures ANOVA (rmANOVA) analysis conducted on RR and HRV parasympathetic indices in the time (RMSSD) and frequency (HF) domains as well as in the non-linear domain (SD1), with DMT as within-subjects factor and Age as a covariate, yielded a significant overall DMT effect (Multivariate test, *F* (4, 16) = 3.028, *p* = 0.049, partial *η*2 = 0.431, power=0.667). Also, univariate within-subjects analysis revealed significant differences between pre-DMT and post-DMT for RMSSD (F(1, 19)=6.020, p=0.024, partial *η*2 = 0.241, power=0.644), HF (F(1, 19)=9.535, p=0.006, partial *η*2 = 0.334, power=0.834) and SD1 (F(1, 19)=6.242, p=0.022, partial *η*2 = 0.247, power=0.659). Conversely, no differences were found between pre-DMT and post-DMT scores for RR values (F(1,19)=0.039, p=0.846).

**Table 2:** RR and HRV-derived parasympathetic and sympathetic indices. In the first column, “pre” and “post” refer to the pre- and post-DMT sessions, respectively.

|  | RR | RMSSD | HF | SD1 | SI |
| --- | --- | --- | --- | --- | --- |
| | mean ± SD | mean ± SD | mean ± SD | mean ± SD | mean ± SD |
| pre | 873.29±156.51 | 14.10±5.55 | 79.71±50.29 | 10.01±3.94 | 18.16±7.16 |
| post | 882.51±124.91 | 17.48±7.94 | 144.76±131.25 | 12.37±5.61 | 18.22±6.62 |

No significant differences (rmANOVA) were found between the pre-DMT and post-DMT scores for the HRV-related sympathetic marker SI (Stress Index) (F(1,19)=1.658,p=0.213, partial *η*2 = 0.080, power=0.193).

Repeated measures ANOVA conducted on SCL mean scores, with DMT as within-subject factor and Age as a covariate, did not yield any significant difference between the pre-DMT (0.502 ± 0.368) and post-DMT (0.498 ± 0.325) scores (F(1,21)=0.598, p=0.448, partial *η*2 = 0.028, power=0.114).

### 3.2. GDS and PSQI results

The mean GDS scores of the group in the pre-DMT session indicated an overall absence of depressive symptoms (mean ± SD, 7.68 ± 4.27), with only eight out of 25 participants reporting scores above the cutoff (cutoff = 9) indicative of mild depression [42]. Similar GDS scores were observed subsequent to the DMT (mean ± SD, 7.34 ± 4.80). Indeed, rmANOVA, with DMT as within-subject factor and Age as a covariate, did not yield any significant difference between the pre-DMT and Post-DMT scores (F(1,23)=0.510, p=0.510, partial *η*2 = 0.019, power=0.098).

The mean PSQI scores of the group in the pre-DMT session (mean ± SD, 5.20 ± 2.83) indicated a negligible deterioration of sleep quality (cutoff = 5) which is consistent with the age-related physiological changes in some sleep parameters (e.g., reduction in total sleep time; increase in night awakenings) [46]. Similar PSQI scores were found in the post-DMT session (5.48 ± 2.93). Indeed, rmANOVA, with DMT as within-subject factor and Age as a covariate, did not yield any significant difference between the pre-DMT and Post-DMT scores (F(1,23)=1.02, p=0.321, partial *η*2 = 0.043, power=0.163). The above results imply a negligible effect of DMT on both the depression scale and the sleep quality of participants.

### 3.3. Correlation analysis

Pearson correlation analysis between Age and parasympathetic indices revealed weak (*r <* 0.3) non significant positive linear correlations between Age and pre-DMT scores (first three lines of Table 3). However, these correlations became much stronger and significant for the post-DMT scores (*r >* 0.54)(last three lines of Table 3). As an example, in Figure 1 the linear associations between Age and the pre- and post-DMT RMSSD (left panel) and SD1 (right panel) scores are shown. A significant increase clearly emerges when comparing pre-DMT and post-DMT associations, and this qualitative observation is quantitatively confirmed by the statistical analyses reported in Table 3.

**Figure 1:**
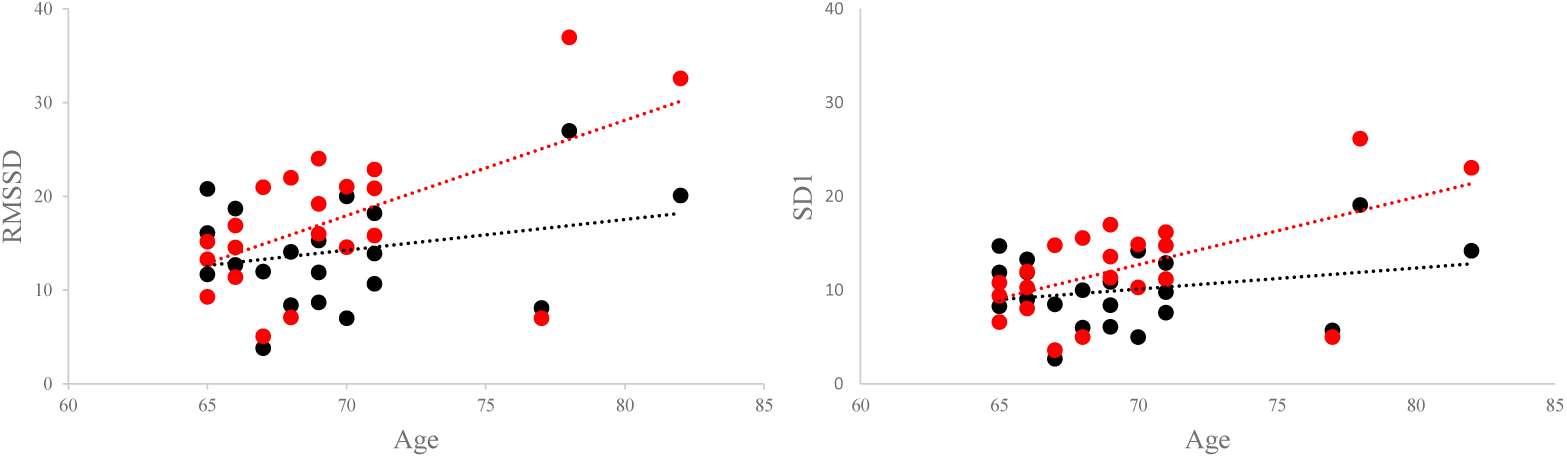
Correlation between Age and pre-DMT (black dots) and post-DMT (red dots) scores of RMSSD and SD1 indices. Trendlines are shown.

**Table 3:** Pearson correlation between Age and parasympathetic HRV indices.

| session | variable | correlation index (r) |
| --- | --- | --- |
| Pre-DMT | RMSSD | 0.266 |
|  | HF | 0.207 |
|  | SD1 | 0.257 |
| Post-DMT | RMSSD | 0.578* |
|  | HF | 0.544* |
|  | SD1 | 0.579* |

Note: \* significance at $p < 0.01$

Pearson’s correlation analysis also showed a negative correlation between RR and SCL mean scores in both sessions (pre-DMT, *r* = −0.483; post-DMT, *r* = −0.421). However, the RR-SCL association was significant only in the pre-DMT session (pre-DMT, *p* = 0.036; post-DMT, *p* = 0.073).

### 3.4. Regression analysis Questionnaires vs. Physiological indices

#### Pre-DMT

The linear regression model indicated that, in the pre-DMT session, GDS scores were predictive of RR (Figure 2). In fact, the model indicated a good significant overall fit (R-square=0.223; F(1,20)=5.465, p=0.030). The regression equation results to be:

**Figure 2:**
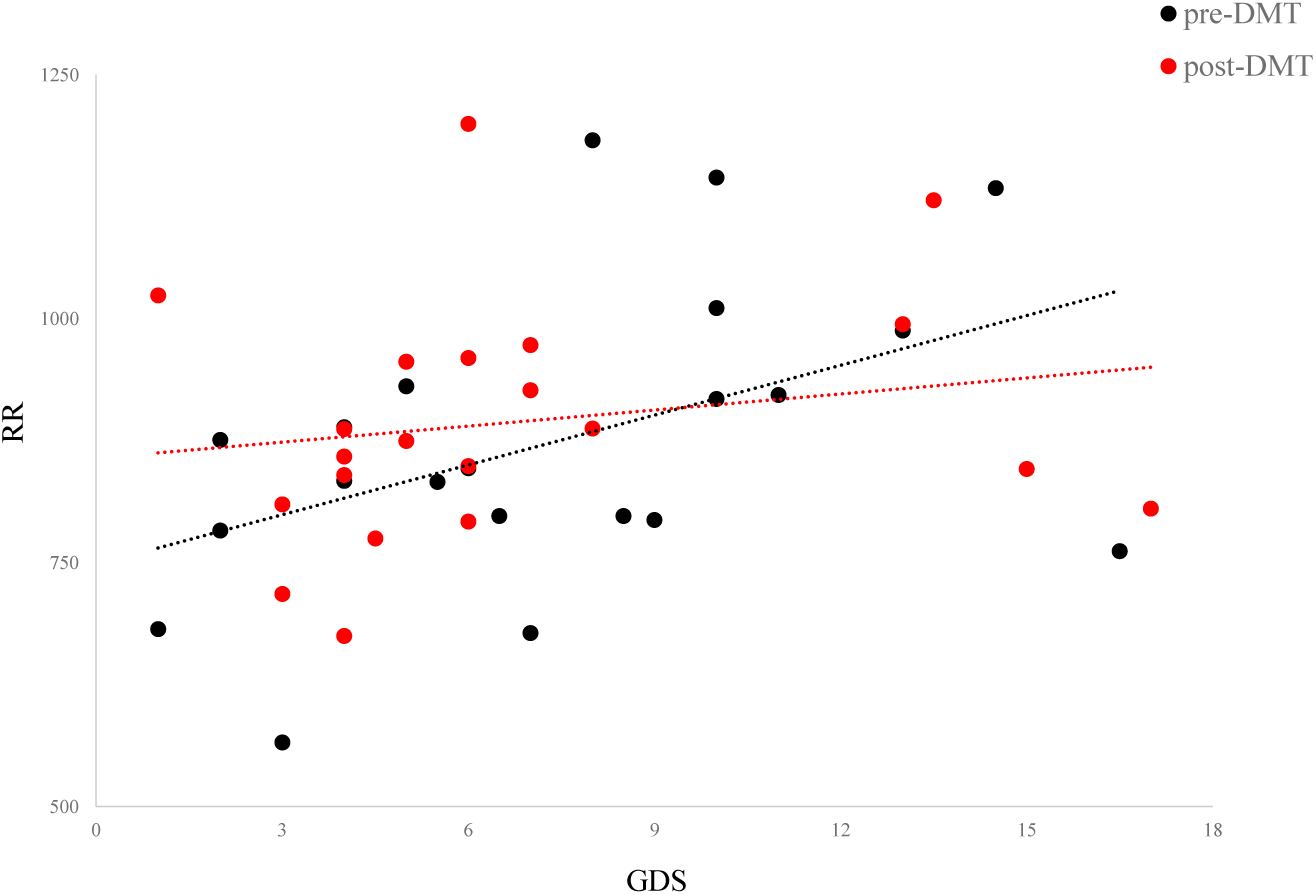
Association between GDS scores and mean RR in the pre-DMT (black dots) and post-DMT (red dots) sessions. Trendlines are shown.

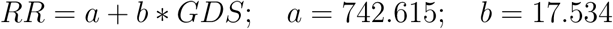

thus indicating that, for each one point increase in the GDS score, RR increases by 17.534 percentage points.

The linear regression results, including the model summary, analysis of variance (ANOVA) and coefficient tables (SPSS), are reported in Figure 1 of the Supplementary Material.

No predictive association was found in the pre-DMT session between GDS scores and HRV indices, GDS scores and SCL as well as between PSQI and RR, HRV indices and SCL.

#### Post-DMT

The association between GDS scores and RR disappeared in the post-DMT session (Figure 2). Predictive associations were still not found in the post-DMT session between GDS scores and HRV indices, GDS scores and SCL as well as between PSQI and all physiological indices (RR, HRV indices and SCL).

## 4. Discussion

The present study is the first one investigating the potential positive effect of DMT on HRV in a sample of healthy older people. The primary finding is that after a 8-week DMT activity a significant increase in resting state HRV parameters (RMSSD, HF, SD1), which are indicators of the parasympathetic outflow to the heart via the vagus nerve, [47] was found. Previous research has shown that high resting state values of RMSSD, HF and SD1, which are known to be positively correlated with each other [2], indicate an active parasympathetic nervous system. The high parasympathetic activity is closely associated with better down-regulation of negative affect, use of adaptive regulatory strategies, more flexible emotional responding [48] and higher cognitive functions (e.g., attention [49] and motor imagery [50]).

This result, that is, the possible efficacy of DMT in improving cardiovascular autonomic regulation in older healthy people, is assessed by post-DMT HRV changes with respect to pre-DMT. This is in agreement with a previous study that employed some form of dance training (i.e., exergaming-based and virtual reality-based dance trainings) as a tool to improve autonomic regulation in the elderly demographic [51, 20]. Nevertheless, the suggested effectiveness of DMT in improving vagally-mediated HRV can not only be attributable to physical activity associated with a pure dance training, but might also be due to the activation of social skills that free body movements stimulate. In fact, the distinctive nature of the DMT approach is the employment of dance and movement to encourage individuals to establish relationships with one another within group dynamics [25, 52, 26, 53].

Interestingly, in the context of social cognition and behaviour, it has been hypothesised that there is a relationship between these domains and vagally-mediated HRV [54, 55, 56]. In fact, HRV has been suggested as a biomarker for adaptive behaviour during social interactions. The existence of this relationship is supported by anatomical and functional data. Indeed, changes in the structure and function of the brain regions involved in social cognition and interactions (prefrontal and paralimbic brain areas) are also associated with changes in vagally-mediated HRV [57]. Also, marked differences in social cognition and social interactions have been found between individuals with different vagal-mediated HRV [58]. Therefore, it can be hypothesised that the increase observed in vagal activity after the participation in DMT activity may be a manifestation of the improvement in social behaviour it facilitated.

Another interesting finding is the strengthening of the (positive) correlation between age and vagal-related HRV indices. In fact, this association becomes much stronger after DMT, thus indicating both a higher vagal control in the eldest participants but also a role of DMT in reinforcing the age-HRV relationship. The higher vagal control in the eldest people appears to contradict previous findings suggesting a decline in HRV with age [7]. However, it is generally accepted that HRV values tend to stabilise after the age of 50 [59]. It is noteworthy that the sample of old people under consideration consists of individuals who participate in leisure motor activities to a moderate extent, such as walking and cycling. They also have a normal BMI, are not affected by any mental or physical disorder and have a fairly good sleep quality. Therefore, given that all these factors are involved in the maintenance of optimal HRV, the observation that vagal indices are higher in the eldest participants underscores the notion that, in this demographic, good health and physical and social activity are effective in reducing age-related vulnerabilities to factors such as stress, anxiety and loneliness, that have the potential to compromise cardiac function.

The SCL-RR negative correlation essentially is found not to change between pre- and post-DMT sessions, even if it is worth noting that only during the pre-DMT session the correlation reached a level of statistical significance. This is not an unexpected outcome. In fact, under conditions of rest, the autonomic balance is dominated by the tonic control of the parasympathetic nervous system, which is associated with a reduced sympathetic drive.

A positive causal association is found between GDS and mean RR. This means that individuals who report a higher number of depressive symptoms display lower heart rates. This finding contrasts with previous research indicating an inverse relationship between GDS and HRV indices, including mean RR [60, 10]. However, it is noteworthy that the mean GDS score fell below the cut-off point for depression, with only a few participants reporting a higher score that indicated very mild depression. Therefore, we can hypothesise that the previously reported negative association emerges exclusively in patients clinically diagnosed with depression. However, it is intriguing to note that administration of DMT disrupts the positive association between GDS and mean RR.

Regarding PSQI, in contrast with previous findings indicating that diminished sleep quality is associated with reduced HRV [35], we find no significant association between PSQI and HRV indices. However, the PSQI scores of both sessions reveal that sleep quality in our sample is relatively good. Only a small number of participants demonstrate scores slightly above the cutoff value, which is consistent with the physiological age-related changes in sleep structure. Therefore, the absence of association between PSQI and HRV and of DMT-related enhancements in the participants’ sleep quality is not unexpected, given that their current sleep quality is already satisfactory.

The results highlighted in this study are encouraging and further validation is required on a larger sample. Actually, the small sample size is one limitation of the study. However, the DMT professional strongly recommended no more than 14 participants per cycle to enable better group dynamics. Furthermore, we put forth that, as already discussed in the Introduction, we applied a one-group pretest–posttest design, thus implementing a well-defined and largely exploited quasi-experimental research method [34]. Further advances in our investigation should involve the use of a control group in order to confirm our preliminary findings. However, this raises questions about which control protocol would be the most appropriate, e.g., structured physical activity, sedentary social activity, or no additional physical or social activity. The final option would probably not be realistic, since participation in our study was dependent on the offer of free participation in an 8-week leisure activity. Additionally, the majority of the participants were specifically attracted by the novelty of the DMT activity.

A further significant issue still related to the complexity of the enrolment process is the fact that, despite the response rate to recruitment advertisements was high, only few people met the eligibility criteria. However, as a countermeasure regarding the lack of a control group, we paid particular attention to any salient personal events or experiences that participants encountered during the eight-week DMT activity (the so called “history effects” [34]), which might have affected the physiological outcome. Indeed, one participants of the Cycle 2, who underwent minor incidents, was excluded from analyses, while none of the other participants reported events outside their daily routine.

Another limitation is the imbalance in the gender of participants, with the majority of the subjects being female. The gender imbalance seen in dance-related activities for older adults has been reported previously, with most studies including a higher percentage of females [17]. The present study attempted to mitigate this imbalance, maintaining a similar female-to-male ratio in both DMT cycles. One final issue is that the study only examined the short-term effects of DMT on HRV and SCL. Future studies may involve taking repeated measurements over short and long periods of time.

In conclusion, the present study provides preliminary evidence that DMT could be an effective strategy for improving the health and wellbeing of older people by preventing or counteracting the age-related decline in HRV that is frequently observed. We argued that this may be ascribed to the DMT peculiarity of stimulating individuals interaction and integration in a collective unit [26, 53]. In this regard, DMT has the potential to promote a shared positive state of mind and facilitate social interactions and wellbeing, also among most senior individuals [33]. Finally, based on these premises, the present study may also be relevant to the underlying framework of embodied cognition [28, 26].

## Supporting information

Explanation of the experimental protocol and Table with details on statistical regression results

## Appendix A. Acronyms and nomemclature

DMT: Dance Movement Therapy
HRV: Heart Rate Variability
ECG: ElectroCardioGram (instrument: g-Nautilus 32 system)
RR: time interval between two R-peak
LF: Low-Frequency component of the power spectrum density of RR (0.04 −0.15 Hz)
HF: High-Frequency component of the power spectrum density of RR (0.15−0.4 Hz)
SD1: Poincaré plot short term variability (in Poincaré plot, the standard deviation perpendicular to the line-of-identity, see [39])
SI: Stress Index (square root of Baevsky’s stress index [41])
RMSSD: Root Mean Square of Successive (RR) Differences
pNN50: the percentage of successive NN (Normal-to-Normal) intervals that differ by more than 50 milliseconds
PSNS: ParaSympathetic Nervous System
SNS: Sympathetic Nervous System
GSR: Galvanic Skin Response (instrument: Shimmer3 GSR+)
SCL: Skin Conductance Tonic Level
ANOVA: ANalysis Of Variance
rmANOVA: repeated measures ANOVA
GDS: Geriatric Depression Scale
PSQI: Pittsburg Quality of Sleep Index

## Funding

This publication was partially supported by European Union - Next Generation EU, in the context of The National Recovery and Resilience Plan, Investment 1.5 Ecosystems of Innovation, Project Tuscany Health Ecosystem (THE), CUP: B83C22003930001, by the University of Pisa and by the Signal and Images Laboratory of ISTI-CNR.

## Acknowledgements

We greatly acknowledge Davide Moroni for valuable discussion and Paolo Orsini for the excellent technical assistance.

## Author contributions statement

L.S.: Conceptualization of experimental protocol; Conceptualization of data analysis; Data collection; Data curation; Formal analysis; Investigation; Methodology; Resources; Supervision; Validation; Visualization; Roles/Writing - original draft; and Writing - review and editing. S.D.: Conceptualization of experimental protocol; Selection criteria and recruitment of participants; Data collection; Investigation; Visualization; Roles/Writing - review and editing. G.I.: Data collection, methodology, GSR data curation, Roles/Writing review and editing. M.R.: Preparation of DMT settings; conduction of DMT sessions while acting as a DMT professional; Data collection; Roles/Writing review and editing. P.P.: Conceptualization of experimental protocol; Preparation of DMT settings; Selection criteria and recruitment of participants; Data collection; Formal analysis; Funding acquisition; Investigation; Methodology; Project administration; Resources; Supervision; Roles/Writing - review and editing.

All authors approved the final version of the paper.

## Statement of Ethics

This study was performed in accordance with the World Medical Association Declaration of Helsinki ethical standards and approved by the Committee on Bioethics of the University of Pisa (resolution no. 29/2024 of 26/07/2024). All participants read and signed the informed consent to participate in the study.

## Competing interests

The authors declare no competing interests.

## Footnotes

1 DMT is defined and regulated by the American Dance Therapy Association (ADTA) [21] and the European Association Dance Movement Therapy (EADMT) [22]. Italian DMT professionals are associated with the Italian Professional Association of Dance-Movement-Therapy (Associazione Professionale Italiana DMT, APID [23]), which also provides three-year training courses with a final thesis/dissertation and a practical test.

2 The sex categories female/male were referred to “sex assigned at birth”.

## Notes

### Competing Interest Statement

The authors have declared no competing interest.

