## Supplementary material for "Exploring Dance Movement Therapy as a Novel Approach to Improving Heart Rate Variability in Healthy Older Adults": Explanation of the experimental protocol and Table with details on statistical regression results

by

Laura Sebastiani, Said Daoudagh, Giacomo Ignesti, Marina Raglianti and Paolo Paradisi

**DMT protocol**

The DMT intervention comprised a series of eight weekly meetings, each with a duration of approximately one hour, held in a standard gymn with ample space for unrestrained movement.

As a DMT professional (conductor) associated with APID (APID 2024), MR conducted the DMT intervention.

The conductor planned the general outline of the settings for each meeting, including suggestions, possible tools - like laces - and the sequence of musical pieces. The same settings were applied in the second cycle of DMT, and the order of meetings was maintained as in the first cycle.

Each meeting began with a 'welcome' phase in which participants were guided by the conductor to take a position within a circle. The aim of this phase was to give participants the opportunity to introduce themselves and to familiarise with the other group members. They achieved this by introducing themselves and then calling each other by name.

The following stage involved a "warm-up" that was accompanied by music. During this phase, participants engaged in preparatory activities for their bodies. These activities included joint articulation and the mobilisation of various anatomical structures.

The focus of each DMT meeting was on a specific thematic element (e.g. exploring the space around the body). The theme was developed in three phases, involving individual, dyadic, and group motor dialogues, with or without accessories (e.g. elastic ribbons, masks, veils). Each phase of the process was accompanied by appropriate music.

The meeting's concluding part generally involved a group activity, which was structured around a soundtrack characterised by pleasant and/or joyful music.

In the context of certain meetings, the conductor suggested implementing a brief 'final greeting' sequence. This involved all participants standing in a circular formation, hand in hand, all together taking two steps to the right and one to the left.

**Regression results from SPSS**

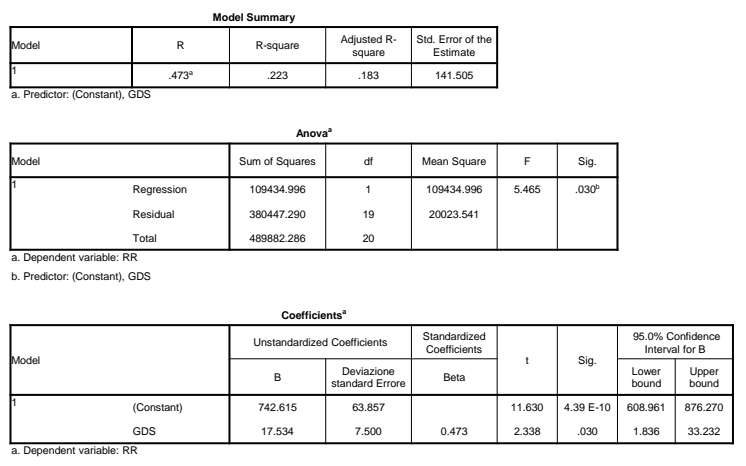
Figure 1: Detailed output of the linear regression analysis carried out with SPSS and applied on mean individual RR vs GDS relative to the pre-DMT session.

**Legenda**

**Model Summary Table**

- ***R*** is the strength of the correlation between our two variables.
- ***R* Square** tells us how much of the variance in the dependent variable is explained by the independent variable.
- **Adjusted** ***R* Square** adjusts ***R* Square** on the basis of the sample size.

**ANOVA Table**

To determine whether our regression model predicts the dependent variable better than we would expect by chance.  p-value <0.05 indicates that the regression model is significant.

**Coefficients Table**

This table gives the values we need to write the regression equation *Y*= *a* + *bX*.

- *a given by column* **B (Constant) - Unstandardized Coefficient:** represents the value of *Y* when *X=0.*
- *B given by column* ***B GDS* - Unstandardized Coefficient:** Slope: the expected change in *Y* for a one-unit increase in *X*.
- **Std. Error - Unstandardized Coefficient:** Standard error of the estimate
- **Beta - Standardized Coefficient:** Shows the expected change in the dependent variable (Y) in standard deviation units.
- **t and Sig.:** Tell whether the independent variable (X) has a significant effect on the dependent variable (Y). If p-value is less than 0.05, the independent variable has a statistically significant effect on the dependent variable.

t = T-statistic ; Sig. = p-value

- **95% Confidence interval for B:** shows that you can be 95% confident that the slope falls in this range.
